# Evolutionary proteomics reveals distinct patterns of complexity and divergence between Lepidopteran sperm morphs

**DOI:** 10.1101/384164

**Authors:** Emma Whittington, Tim Karr, Andrew J. Mongue, Steve Dorus, James R. Walters

## Abstract

Spermatozoa are one of the most strikingly diverse animal cell types. One poorly understood example of this diversity is sperm heteromorphism, where males produce multiple distinct morphs of sperm in a single ejaculate. Typically, only one morph is capable of fertilization and the function of the non-fertilizing morph, called parasperm, remains to be elucidated. Sperm heteromorphism has multiple independent origins, including Lepidoptera (moths and butterflies), where males produce a fertilizing *eupyrene* sperm and an *apyrene* parasperm, which lacks a nucleus and nuclear DNA. Here we report a comparative proteomic analysis of eupyrene and apyrene sperm between two distantly related lepidopteran species, the monarch butterfly (*Danaus plexippus*) and Carolina sphinx month (*Manduca sexta*). In both species, we identified approximatey 700 sperm proteins, with half present in both morphs and the majority of the remainder specific to eupyrene sperm. Apyrene sperm thus have a distinctly less complex proteome. Gene Ontology (GO) analysis revealed proteins shared between morphs tend to be associated with canonical sperm cell structures (e.g. flagellum) and metabolism (*e.g*. ATP production). GO terms for morph-specific proteins broadly reflect known structural differences, but also suggest a role for apyrene sperm in modulating female neurobiology. Comparative analysis indicates that proteins shared between morphs are most conserved between species as components of sperm, while morph-specific proteins turn over more quickly, especially in apyrene sperm. The rapid divergence of apyrene sperm content is consistent with a relaxation of selective constraints associated with fertilization and karyogamy. On the other hand, parasperm exhibit greater evolutionary lability, which may reflect adaptive response to shifting regimes of sexual selection. Additionally, we provide the first (to our knowledge) scanning electron micrographs of lepidopteran sperm.

## Introduction: The enigma of sperm heteromorphism

Sperm heteromorphism is a phenomenon in which males produce multiple distinct sperm morphs as a developmentally normal and regulated process during gametogenesis. This phenomenon has arisen across a broad range of taxa, including multiple independent origins in Insecta and Mollusca, and a few vertebrates (Swallow and Wilkinson 2002; Till-Bottraud et al. 2005; Hayakawa 2007). Sperm morphs are defined by their fertilization capacity, with only one morph (eusperm) capable of successful fertilization, whereas the remaining morphs are not (parasperm) (Healy & Jamieson 1981). Although parasperm function has yet to be conclusively determined in any taxa, two adaptive hypotheses have been investigated: 1) facilitation of eusperm or 2) mediation of sperm competition (reviewed in Swallow & Wilkinson 2002; Till-Bottraud et al. 2005). The former hypothesis is supported by observations that fertility can be strongly impacted by parasperm absence or variety (Sahara & Kawamura 2002; Sahara & Takemura 2003; Oppliger et al. 2003). The latter is supported by the tailoring of parasperm investment in response to the intensity of sperm competition (He & Miyata 1997; Oppliger et al. 1998; Wedell & Cook 1999). As would be expected given their presumed functional diversification, parasperm and eusperm exhibit distinct evolutionary patterns, with parasperm diverging faster and showing greater predicted evolvability (Moore et al. 2013; Snook 2017; Holman et al. 2008).

One striking example of sperm heteromorphism occurs in Lepidoptera (moths and butterflies), where the parasperm morph, called apyrene sperm, lacks a nucleus and nuclear DNA (thoroughly reviewed by Freidlander and Reynold (2005)). Apyrene sperm are present in all studied species of Lepidoptera except for the most ancestrally-diverging lineage. Thus, sperm heteromorphism appears to have arisen ancestrally and has been retained across taxa with diverse mating systems and sperm competition intensities. In most species, apyrene sperm vastly outnumber their nucleated eusperm counterparts (called *eupyrene* sperm), typically accounting for ~85-90% of sperm produced. Numerous microscopy studies contrasting apyrene and eupyrene sperm have revealed several structural differences beyond apyrene sperm lacking a nucleus, including: 1) apyrene sperm lack an acrosome, 2) apyrene sperm are shorter, and 3) prior to ejaculation, eupyrene sperm remain bundled in an extracellular sheath (Fig. 1). However, very little is known about how these sperm morphs differ at the molecular level.

**Figure 1.**
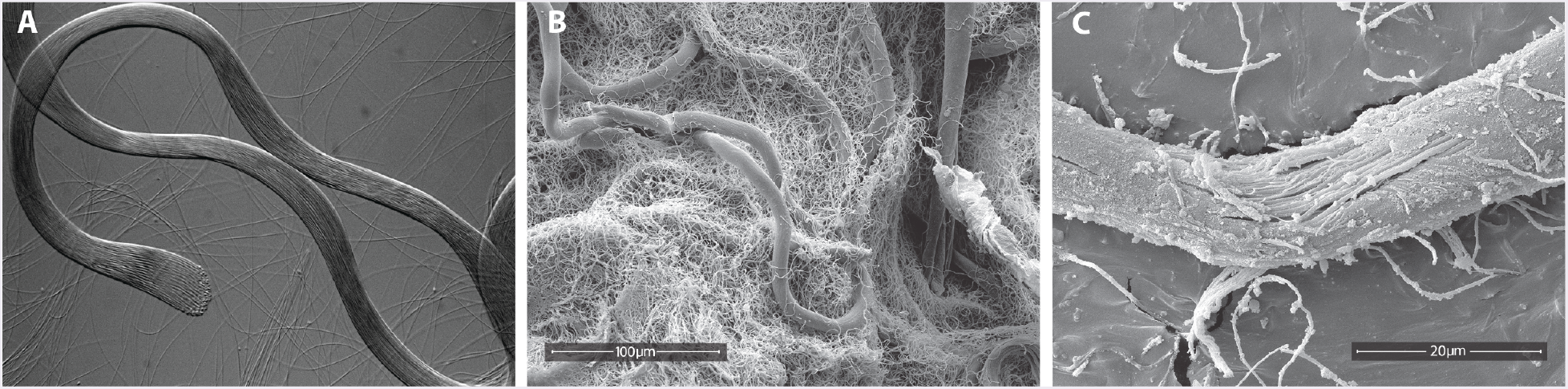
Microscopy of *Manduca* eupyrene sperm bundles and apyrene sperm dissected from the male reproductive tract. (A) Differential interference contrast image of a single bundle of eupyrene sperm, with unbundled individual apyrene sperm visible in the background. (B) Scanning Electron Microscopy (SEM) image of sheathed eupyrene bundles amongst an abundance of apyrene sperm. (C) SEM image of a eupyrene sperm bundle with a section of the sheath-matrix removed to reveal sperm tails. To our knowledge, these are the first SEM images of lepidopteran sperm in the public record; additional SEM images are provided in Supplemental file 2.

In previous studies, we employed high-throughput liquid chromatography tandem mass spectrometry (LC-MS/MS) proteomics to analyze co-mixed apyrene and eupyrene samples from both and the Carolina sphinx moth (*Manduca sexta*, henceforth *Manduca*) and monarch butterfly (*Danaus plexippus*) (Whittington et al. 2015; 2017). These studies revealed substantial divergence in proteome content between species but did not included direct comparisons between morphs. However, morph-specific proteome analysis using two-dimensional gel electrophoresis in monarch indicated that the eupyrene sperm proteome was notably more complex (Karr & Walters 2015). Extending analyses to separately characterize apyrene and eupyrene sperm proteomes, respectively, will establish their molecular differences and potentially inform our understanding of apyrene sperm function.

Here we report the results of LC-MS/MS proteomic analysis applied to isolated apyrene and eupyrene sperm samples. Doing so in both monarch and *Manduca* allowed us to contrast the overlap in protein content and function between morphs and between species. We found proteome composition has diverged more rapidly for morph-specific than shared proteins. Functional annotations broadly reflect known structural differences, and hint at a role for apyrene sperm in modulating female neurobiology.

## Results I: Proteome Composition, Complexity, and Function

LC-MS/MS analysis of isolated apyrene and eupyrene sperm samples identified a combined total of 742 proteins in *Manduca* and 661 proteins in monarch (Fig. 2; supplementary tables S1 & S2). Our analysis confirms the previous observation in monarch that the apyrene sperm is less complex in protein composition relative to eupyrene sperm (Karr & Walters 2015), and extends this observation to *Manduca*. Also, in both species, approximately half of identified proteins were shared between sperm types (Fig. 2). Given that these two species diverged over 100 mya (Heikkilä et al. 2012), it seems likely that reduced apyrene proteome complexity and substantial overlap between morphs is generally representative of Lepidoptera. Nonetheless, the proportions of proteins unique to or shared between morphs differed substantially between species (Fig. 2; c^2^ = 47.1, d.f. = 2, p < 0.0001), primarily reflecting monarch’s greater disparity in complexity between morph-specific proteomes.

**Figure 2.**
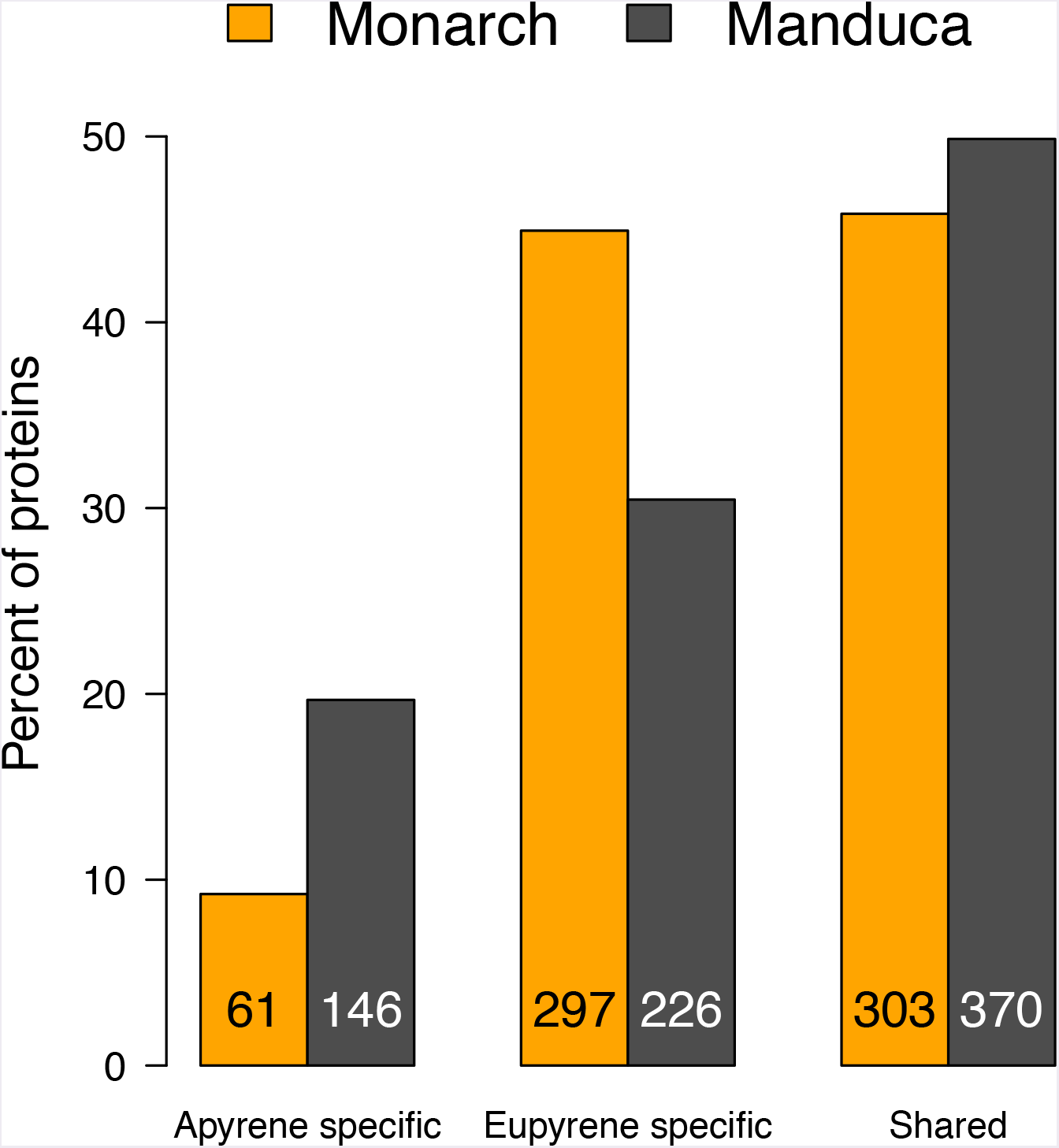
Portion of proteins found in each subset of the sperm proteome. Proteins identified only in apyrene or eupyrene sperm are “specific”, while proteins identified in both morphs are “shared”. Bar heights represent percent of total proteins. Numbers at the base of bars are the counts of proteins identified in each subset.

In both species, approximately three-quarters of sperm proteins were successfully annotated with Gene Ontology (GO) terms, with no significant difference in the proportion of annotated proteins between the morph-specific or shared subsets (c^2^ < 4.2 in both species, d.f. = 1, p > 0.05). GO-term enrichments highlight broad functional distinctions among morph-specific and shared proteins (Table 1; electronic supplementary material S3). While apyrene and eupyrene sperm necessarily play distinct (though yet unresolved) roles in fertilization, these discrete morphs have many similarities in morphology (e.g. axonemal based flagellum), physiology (e.g. ATP production) and behavior (e.g. motility) (Friedlander et al. 2005). Thus shared proteins are expected to be enriched for GO-terms related to these functions, as well as others also associated with spermatozoa. Consistent with this prediction, in both species the shared set of proteins tend to have broad associations with the cytoskeletal structure, mitochondria, and cilia (*N.B*. the sperm flagellum is a modified cilium (Dallai 2014)).

**Table 1.**
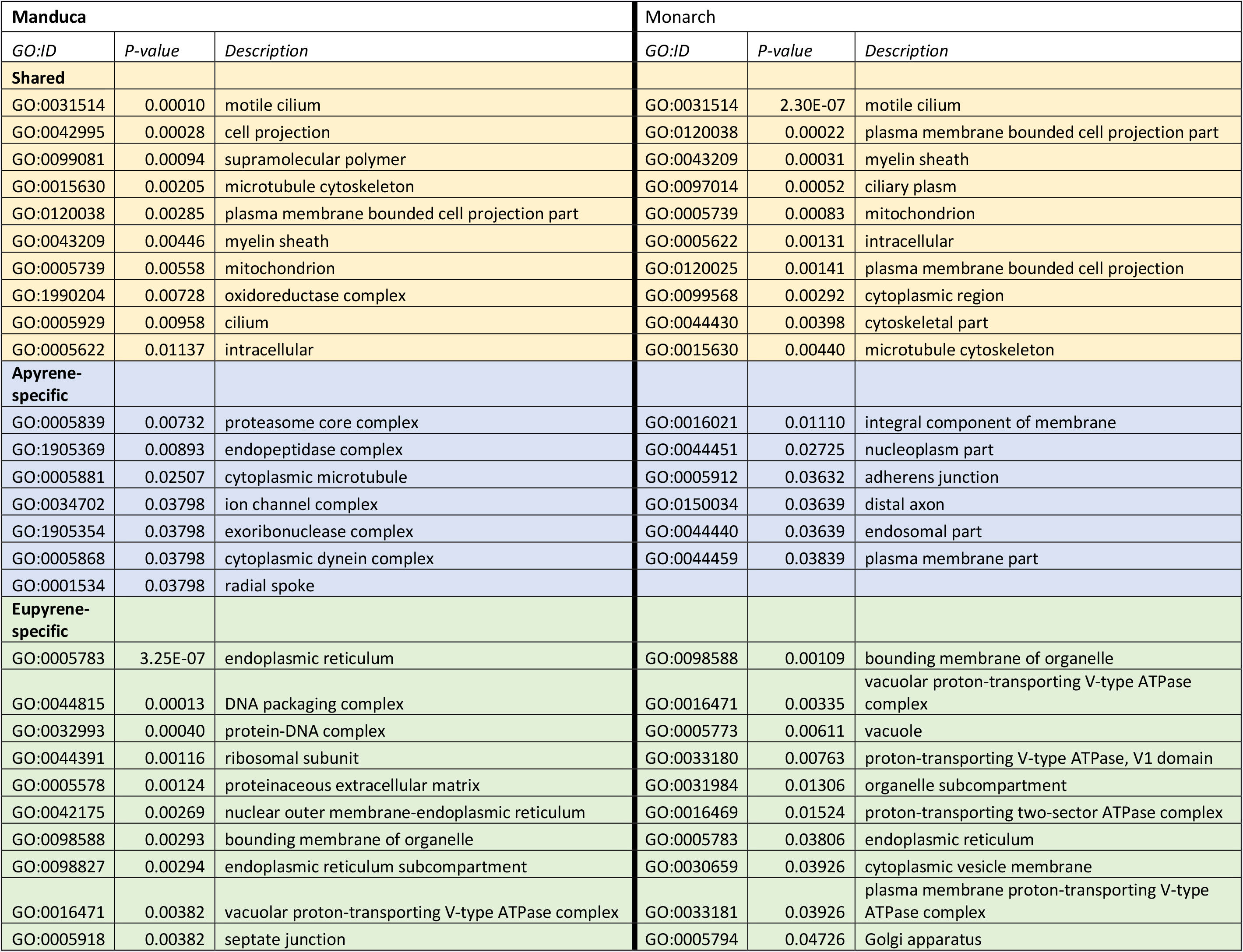
Gene Ontology (GO) Cellular Component terms significantly enriched in subsets of lepidopteran sperm.?

Similarly, structural differences between apyrene and eupyrene sperm are observed in morph-specific GO-term enrichments. Most prominently, apyrene sperm lack a nucleus and nuclear DNA and the endoplasmic reticulum (ER) is greatly reduced during development and presumably absent in mature apyrene sperm (Wolf 1992). Accordingly, among eupyrene-specific proteins, GO-terms associated with the ER and nuclear membrane are enriched, particularly in *Manduca*. Also, in *Manduca*, terms associated with protein-DNA packaging are among the most significantly enriched in eupyrene-specific proteins, reflecting chromatin-related proteins absent from apyrene sperm. Apyrene sperm also lack an acrosome, a vesicle/vacuole organelle typically located in the head of sperm; a corresponding enrichment for vacuole-related GO-terms was identified among eupyrene-specific proteins in both species. Finally, an enrichment of terms associated with extracellular structures is unique to eupyrene-specific proteins. In contrast to apyrene sperm, eupyrene sperm from an individual cyst are packaged and transferred to the female in bundles, sheathed in a proteinaceous extracellular matrix that is subsequently degraded in the female (Fig. 1; additional SEM figures are given in Supplemental file 2). Eupyrene sperm samples in this study were isolated from seminal vesicles and thus were still bundled, hence these “extracellular” eupyrene proteins likely comprise this sheathing.

While GO-term analysis at the molecular level broadly mirrors previously known structural differences between sperm morphs, it does not yield much insight into apyrene sperm function. We find few commonalities between species among GO-terms enriched in apyrene-specific proteins (electronic supplementary material S3) and this may reflect distinct apyrene functions in our study species. Parasperm potentially act as vehicles transporting molecules to the female reproductive tract, as is proposed in some mollusk species (Lobov et al. 2018). Along these lines, *neuron development* is the most significantly enriched “Biological Process” term among monarch apyrene-specific proteins. Other terms associated with neuronal-development are also enriched in *Manduca* (though with less significance; 3). It is well-known in *Drosophila* that components of the male ejaculate impact female neurobiology, modulating post-mating shifts in behavior and physiology (Chow et al. 2013). In *Helicoverpa armigera* moths, male accessory gland extracts produce a strong post-mating response in females (Fan et al. 1999), mediated by a receptor specifically expressed in both female neural and reproductive structures (Hanin et al. 2011; 2012). Thus, there is precedent for male-derived proteins to modulate female neuro-endocrinology and reproductive physiology. It is therefore plausible that apyrene sperm deliver neuro-endocrine active proteins that modulate female postmating responses.

## Results II: Sperm protein orthology and homology

Spermatozoan morphology is strikingly diverse across animals, an observation seemingly at odds with their fundamental role in reproduction, which should arguably result in evolutionary constraint. This diversity is often explained as the outcome of sexual selection, particularly sperm competition, which can drive the rapid evolution of male reproductive characters (Pitnick et al. 2009). It has been suggested that non-fertilizing parasperm allow the resolution of these potentially conflicting selective pressures via division of labor. For instance, eusperm may shoulder the selective constraints of fertilization, while parasperm primarily function in sperm competition and experience reduced evolutionary constraint and increased adaptive selection (Kura & Nakashima 2000). Consistent with this, quantitative genetic analyses indicate greater evolvability in *Drosophila pseudoobscura* parasperm (Moore et al. 2013). Together, these hypothesis offers a plausible explanation for the contrasting patterns of gene orthology versus sperm homology we observe among lepidopteran sperm morphs (Fig. 3). In the genome of each species, proportions of identifiable orthologs are indistinguishable across the three subsets of proteins (Fig. 3, black). However, the pattern is strikingly different for rates of *sperm homology* (orthologs that are present in the sperm of both species, regardless of morph; Fig. 3, blue). Sperm homology is greatly reduced for morph-specific proteins relative to orthology. For shared proteins, this effect is much less pronounced. Thus, proteins shared between sperm morphs are also relatively conserved as sperm proteins across species, likely reflecting morphological and physiological characteristics broadly common to sperm. In contrast, morph-specific protein content turns over more rapidly than gene gain/loss occurring between species at the whole genome level.

**Figure 3.**
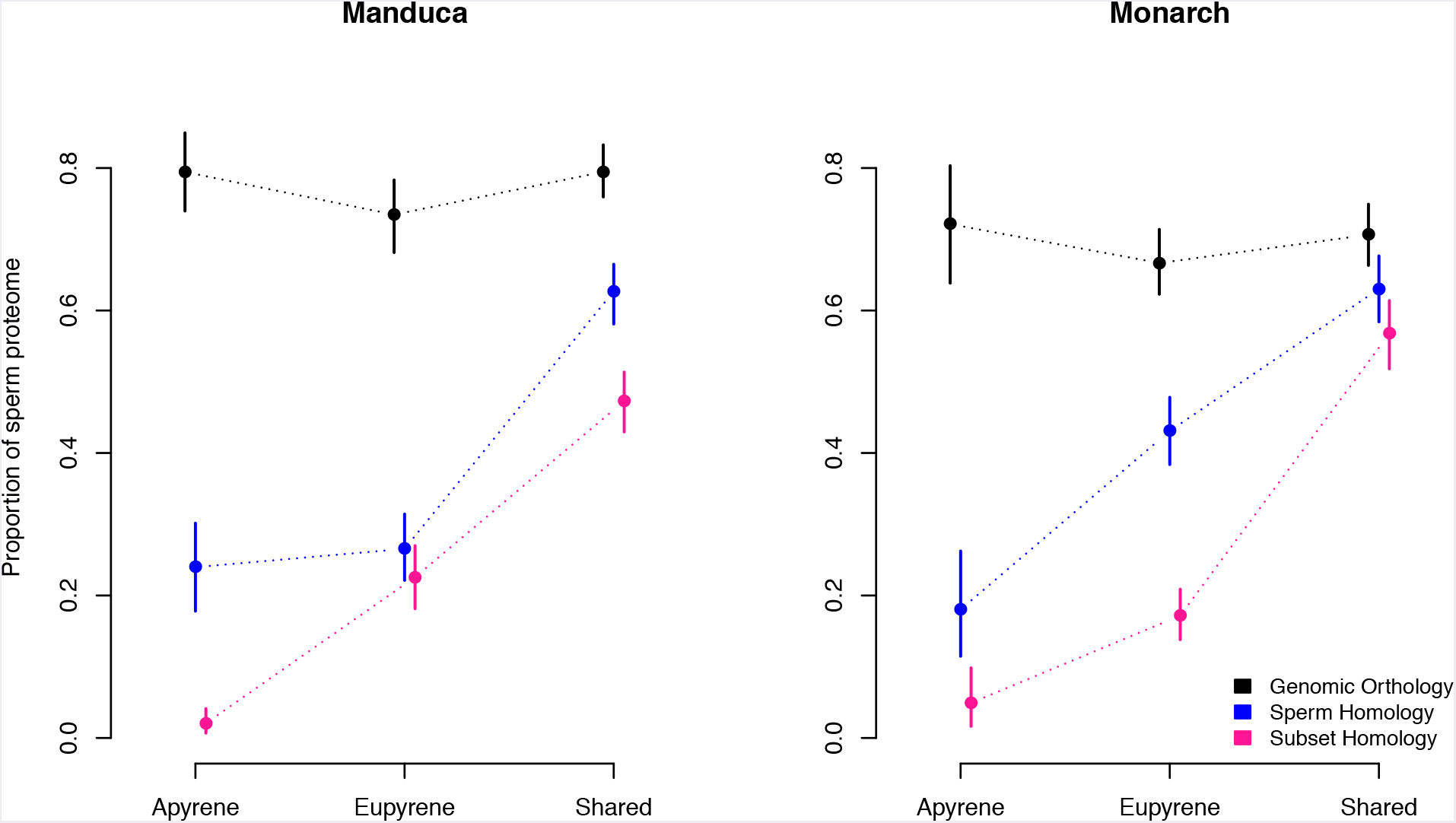
Proportions of proteins homologous between monarch and *Manduca* in each subset of the sperm proteome. The proportion of homologs are plotted for each subset of the sperm proteome: apyrene-specific, eupyrene-specific, and shared. Three different, increasingly stringent, criteria for homology are displayed. Genomic orthologs (black) are proteins found in the genomes of both species. Sperm homologs (blue) are genomic orthologs found in the sperm of both species, regardless of sperm morph. Subset homologs (magenta) are sperm homologs found in the same subset of the sperm proteome.

Comparing morph-specific subsets, our results suggest protein turn-over is faster in apyrene than eupyrene sperm. This is most prominent in monarch, where sperm homology is significantly lower among apyrene-specific than eupyrene-specific proteins (based on nonoverlapping 95% confidence intervals). This difference is not apparent in *Manduca* until examining *subset homology* (requiring orthologs to be found in the same subset of the sperm proteome in both species; Fig. 3, magenta). Subset homology shows similar patterns between species and strongly indicates that apyrene-unique proteins are the least conserved among the subsets examined. Only three orthologs were found to be unique to apyrene sperm in both species.

## Discussion: Apyrene sperm evolution

The relatively rapid turnover of the apyrene-specific proteome is consistent with parasperm experiencing distinct selective pressures compared to eusperm. It is unclear whether this reflects relaxed constraint, greater adaptation, or some combination thereof. As the nonfertilizing sperm morph, apyrene sperm are free of constraints associated with egg interactions, karyogamy, and embryogenesis (Snook & Karr 1998). Consequently, proteins involved in these processes become superfluous to apyrene sperm function. The reduced complexity of the apyrene sperm proteome therefore likely reflects streamlining of a eupyrene “ancestor” present at the root of Lepidoptera. This process might be expected to be random, causing differential protein loss between lineages. Additionally, freed from selective constraints apyrene sperm may undergo lineage specific and adaptive functional specialization, thus compounding the pattern of increased divergence among apyrene-specific proteins. Mating system and the intensity of sperm competition are likely to affect the rate and direction of specialization. Notably, females are far more promiscuous in monarch than *Manduca* (Snow et al. 1974; Drummond 1984), which may explain some of the differences between species observed here. Discerning the relative contributions of relaxed constraint and adaptation to the rapid turnover of apyrene-specific proteins presented here is a novel goal for sperm heteromorphism research. In generating a substantial list of morph-specific proteins for further research, our LC-MS/MS analysis of lepidopteran sperm represents a major advance towards better understanding the still-enigmatic role of apyrene sperm, and parasperm more broadly.

## Acknowledgements

We thank Monarch Watch for support in rearing monarch butterflies, Sheri Skerget for expert technical assistance, Desiree Forsythe for preliminary analyses and the University of Cambridge Proteomics Facility. Kirsten Jensen and Kaylee Herzog generously assisted with SEM imaging. Computing for this project was performed on the Syracuse University Crush Virtual Research Cloud. Funding for this research was provided by Syracuse University to SD, by University of Kansas (KU) to JRW, by the KU Gould Fellowship to AJM, and by Syracuse University and Marilyn Kerr Fellowships to EW.

## Methods

### Sperm samples and proteomic analysis

Two species were used in this study, the monarch butterfly (*Danaus plexippus*, Monarch Watch, Lawrence, Kansas), and the Carolina sphinx moth (*Manduca sexta*, Carolina Biological, Burlington, NC). Sperm samples were isolated from male seminal vesicles 5 to 10 days post eclosion via a small incision in the mid to distal region of the seminal vesicle. Apyrene and eupyrene sperm were separated and purified as described in Karr and Walters (2015). Samples from 3-5 males were pooled for each of 3 biological replicates in each species, resulting in a total of 12 samples.

Proteomic analyses followed the protocol reported in Whittington *et al*. (2017).

Described briefly, each sample was size-separated on a poly-acrylamide gel and cut into four slices which were individually analyzed via LC-MS/MS. Resulting mass spectra were matched to predicted proteins from each species’ genome using the Trans-Proteomic Pipeline (Zhan and Reppert 2012; Deutsch et al. 2015; Kanost et al. 2016). Proteins included in the final sperm proteomes met the following criteria: (1) identification in two or more biological replicates or (2) identification in a single replicate by two or more unique peptides. Comparing apyrene and eupyrene proteomes results in three data subsets: proteins identified uniquely in either apyrene or eupyrene sperm (i.e. morph-specific proteins), and proteins shared between sperm morphs.

All LC-MS/MS data were deposited to the ProteomeXchange Consortium via the PRIDE partner repository with the dataset identifier PXD010168 (Vizcamo et al. 2016).

### Functional annotation and homology

Functional annotations and Gene Ontology (GO) assignments were generated by PANNZER (Koskinen et al. 2015; Ashburner et al. 2000). GO-term enrichment tests were performed using the GOstats Bioconductor package, employing the “conditional = TRUE” setting to account for hierarchical redundancy in GO classifications (Falcon & Gentleman 2007). Hypergeometric tests used the union of apyrene and eupyrene proteins as the background “universe” of genes to identify terms enriched in morph-specific proteins or proteins shared between morphs.

Orthologs between monarch and *Manduca* gene sets were predicted via the proteinortho pipeline (Lechner et al. 2011) with default settings, using the longest isoform per gene. Orthologous proteins identified in the sperm proteome of both species were further classified as *sperm-homologs*. In a few cases, paralogy resulted in small gene groups with a one-to-many or many-to-many relationship between species. Sperm proteins identified within such paralagous groups in both species were also classified as a sperm-homolog. Proportions of orthology and sperm-homology were compared between the three subsets of proteins, with significant differences assessed as non-overlapping 95% confidence intervals, generated by 1000 bootstrap-replicates.

### Microscopy

For light microscopy, sperm were dissected from Manduca seminal vesicles and imaged using differential interference contrast on an Olympus BX60 microscope.

For SEM images, an adult male *Manduca* was dissected to obtain intact seminal vesicles containing eupyrene and apyrene sperm. This tissue was fixed in 10% formalin for 3.5 hours, and then washed 4 times with 70% ethanol, with 5 minutes in between each wash. Samples were rested for 1 hour and washed twice with distilled water with 30 minutes in between washes. All liquid was removed, and samples were soaked overnight in 1% OsO4 in the dark. The next day, samples were washed twice with distilled water then dried with a series of increasing ethanol washes (70%, 95%, 100%) with 10 minutes rest in between each wash. Seminal vesicles were removed from ethanol, placed on a depressed slide, and treated with hexamethyldisilazine (HMDS) as a final drying step. After 30 minutes, excess liquid was removed, and the samples were air dried for an additional 10 minutes before being placed on a prepared specimen mount stub. After mounting to the stub, seminal vesicles were ruptured with a needle and the contents were spread across the stub using a dissecting pin. The stub was then sputter coated with 35nm of gold and imaged on an FEI Versa 3D Dual Beam microscope at the University of Kansas’s Microscopy and Analytical Imaging Lab.

## Supplementary Online Materials

Supplementary file 1: A-E_SupplemntalTables.xls provides the following supplementary data:

Table S1. Complete list of monarch sperm proteins identified in apyrene or eupyrene sperm, with predicted orthologs and functional description.

Table S2. Complete list of *Manduca* sperm proteins identified in apyrene or eupyrene sperm, with predicted orthologs and functional description.

Table S3. Complete list of enriched Gene Ontology terms for morph-specific and shared proteins in monarch. 3. Complete list of enriched Gene Ontology terms for morph-specific and shared proteins in *Manduca*.

Supplementary file 2: sperm_SEM_captions.pdf provides several additional scanning electron microscopy images of sperm from *Manduca* and monarch.

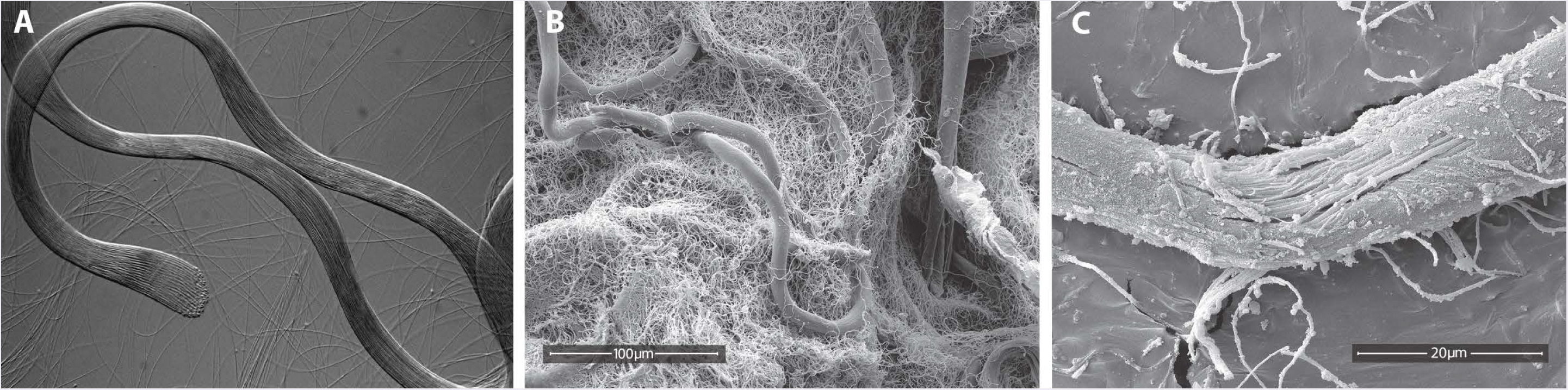

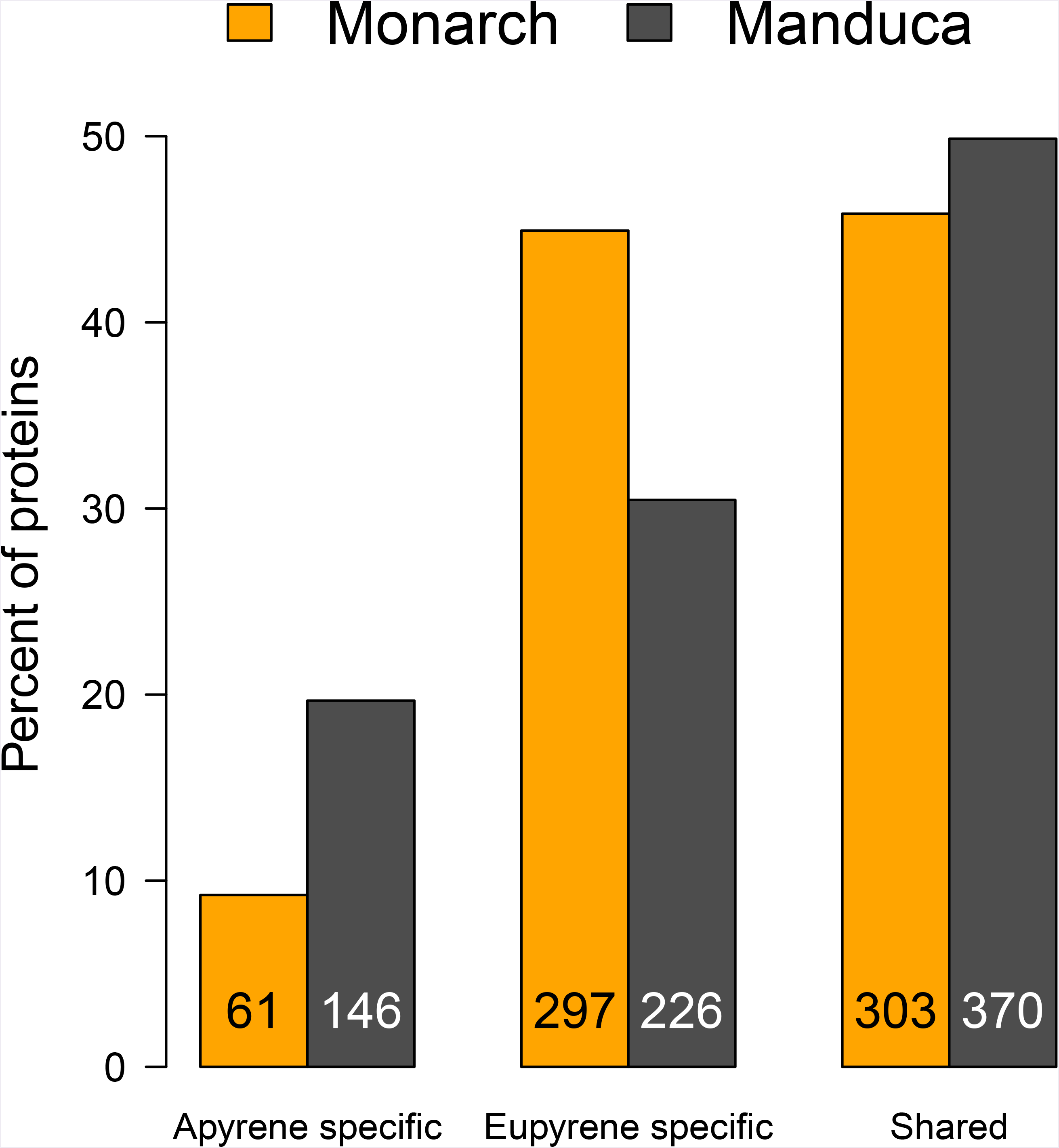

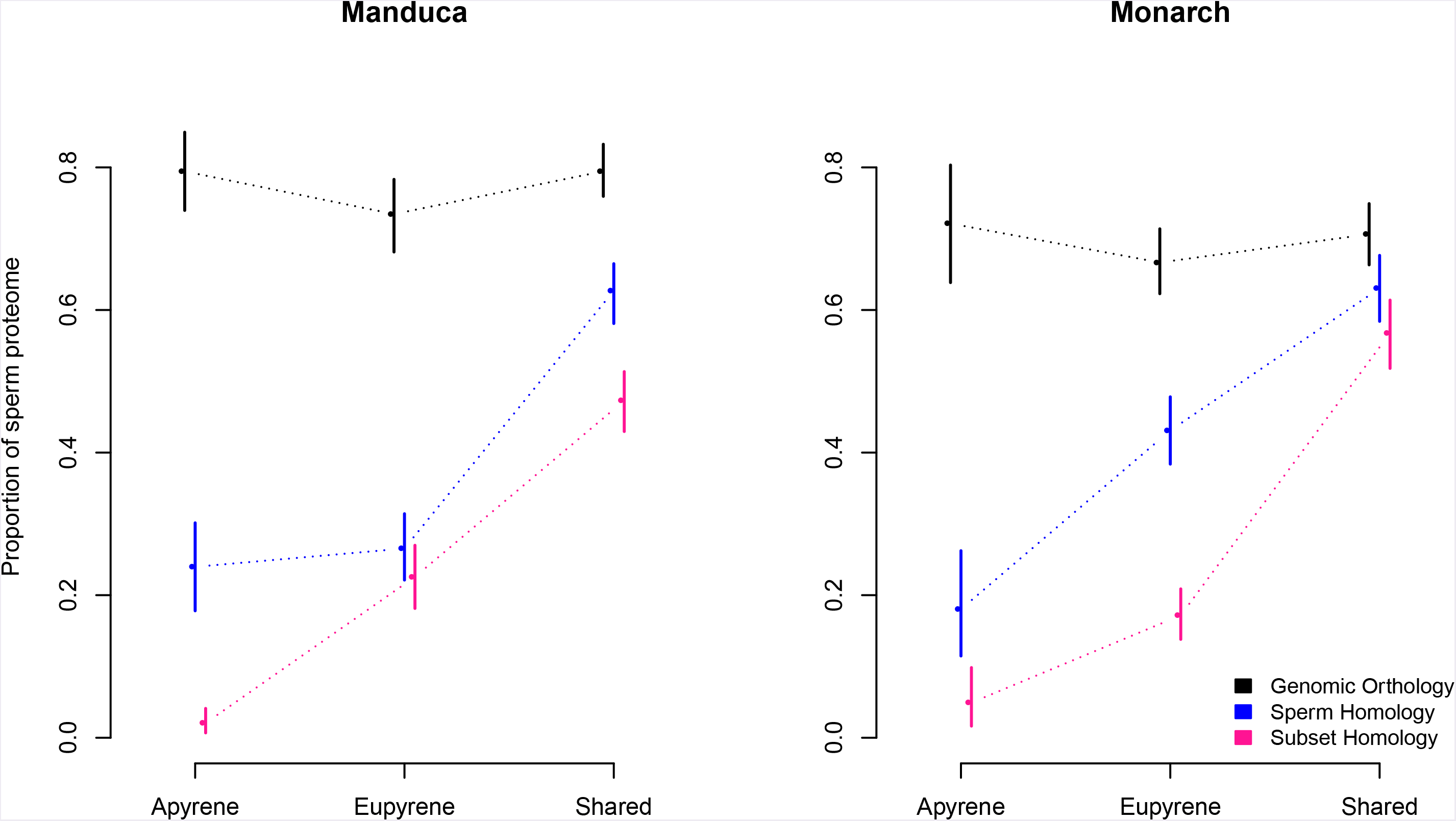

## References

Ashburner M et al. 2000. Gene ontology: tool for the unification of biology. The Gene Ontology Consortium. Nat Genet. 25:25–29. doi: 10.1038/75556.

Chow CY, Wolfner MF, Clark AG. 2013. Large neurological component to genetic differences underlying biased sperm use in Drosophila. Genetics. 193:177–185. doi: 10.1534/genetics.112.146357.

Dallai R. 2014. Overview on spermatogenesis and sperm structure of Hexapoda. Arthropod Structure and Development. 43:257–290. doi:10.1016/j.asd.2014.04.002.

Drummond BA. 1984. Multiple mating and sperm competition in the Lepidoptera. In: Sperm Competition and the Evolution of Animal Mating Systems. Smith, RL, editor. Academic press: New York pp. 291–370.

Falcon S, Gentleman R. 2007. Using GOstats to test gene lists for GO term association. Bioinformatics. 23:257–258. doi: 10.1093/bioinformatics/btl567.

Fan Y, Rafaeli A, Gileadi C, Kubli E, Applebaum SW. 1999. Drosophila melanogaster sex peptide stimulates juvenile hormone synthesis and depresses sex pheromone production in Helicoverpa armigera. Journal of Insect Physiology. 45:127–133.

Friedlander M, Seth R, Reynolds S. 2005. Eupyrene and apyrene sperm: Dichotomous spermatogenesis in Lepidoptera. Adv Insect Physiol. 32:206–308. doi: 10.1016/S0065-2806(05)32003-0.

Hanin O, Azrielli A, Applebaum SW, Rafaeli A. 2012. Functional impact of silencing the Helicoverpa armigera sex-peptide receptor on female reproductive behaviour. Insect Mol Biol. 21:161–167. doi: 10.1111/j.1365-2583.2011.01122.x.

Hanin O, Azrielli A, Zakin V, Applebaum S, Rafaeli A. 2011. Identification and differential expression of a sex-peptide receptor in Helicoverpa armigera. Insect Biochem Mol Biol. 41:537–544. doi: 10.1016/j.ibmb.2011.03.004.

He Y, Miyata T. 1997. Variations in sperm number in relation to larval crowding and spermatophore size in the armyworm, Pseudaletia separata. Ecological Entomology. 22:41–46. doi: 10.1046/j.1365-2311.1997.00030.x.

Healy JM, Jamieson BGM. 1981. An ultrastructural examination of developing and mature paraspermatozoa in Pyrazus ebeninus (Mollusca, Gastropoda, Potamididae). Zoomorphology. 98:101–119. doi: 10.1007/BF00310431.

Heikkila M, Kaila L, Mutanen M, Peña C, Wahlberg N. 2012. Cretaceous origin and repeated tertiary diversification of the redefined butterflies. Proceedings of the Royal Society B: Biological Sciences. 279:1093–1099. doi: 10.1098/rspb.2011.1430.

Holman L, Freckleton RP, Snook RR. 2008. What use is an infertile sperm? A comparative study of sperm-heteromorphic Drosophila. Evolution. 62:374–385. doi: 10.1111/j.1558-5646.2007.00280.x.

Karr TL, Walters JR. 2015. Panning for sperm gold: Isolation and purification of apyrene and eupyrene sperm from lepidopterans. Insect Biochem Mol Biol. 63:152–158. doi: 10.1016/j.ibmb.2015.06.007.

Koskinen P, Törönen P, Nokso-Koivisto J, Holm L. 2015. PANNZER: high-throughput functional annotation of uncharacterized proteins in an error-prone environment. 31:1544–1552. doi: 10.1093/bioinformatics/btu851.

Kura T, Nakashima Y. 2000. Conditions for the evolution of soldier sperm classes. Evolution. 54:72–80.

Lechner M et al. 2011. Proteinortho: Detection of (Co-)orthologs in large-scale analysis. BMC Bioinformatics. 12:124. doi: 10.1186/1471-2105-12-124.

Lobov AA et al. 2018. LOSP: A putative marker of parasperm lineage in male reproductive system of the prosobranch mollusk Littorina obtusata. J. Exp. Zool. (Mol. Dev. Evol.). 330:193–201. doi: 10.1002/jez.b.22803.

Moore AJ, Bacigalupe LD, Snook RR. 2013. Integrated and independent evolution of heteromorphic sperm types. Proceedings of the Royal Society B: Biological Sciences. 280:20131647. doi: 10.1098/rspb.2013.1647.

Oppliger A, Hosken DJ, Ribi G. 1998. Snail sperm production characteristics vary with sperm competition risk. Proc Biol Sci. 265:1527–1534. doi: 10.1098/rspb.1998.0468.

Pitnick SS, Hosken DJ, Birkhead TR. 2009. Sperm morphological diversity. In: Sperm Biology. Sperm Biology: An Evolutionary Perspective Academic Press pp. 69–149.

Snook RR. 2017. Is the Production of Multiple Sperm Types Adaptive? Evolution. 51:797–808. doi: 10.1111/j.1558-5646.1997.tb03662.x.

Snook RR, Karr TL. 1998. Only long sperm are fertilization-competent in six sperm-heteromorphic Drosophila species. 8:291–294.

Snow JW, Copeland WW, Goodenough JL. 1974. The tobacco hornworm: notes on morphology and mating habits. Journal of the Georgia Entomological Society. 9:36–41.

Swallow JG, Wilkinson GS. 2002. The long and short of sperm polymorphisms in insects. Biol Rev. 77:153–182.

Till-Bottraud I, Joly D, Lachaise D, Snook RR. 2005. Pollen and sperm heteromorphism: convergence across kingdoms? Journal of Evolutionary Biology. 18:1–18. doi: 10.1111/j.1420-9101.2004.00789.x.

Vizcamo JA et al. 2016. 2016 update of the PRIDE database and its related tools. Nucleic Acids Research. 44:D447–D456. doi: 10.1093/nar/gkv1145.

Wedell N, Cook PA. 1999. Butterflies tailor their ejaculate in response to sperm competition risk and intensity. Proceedings of the Royal Society B: Biological Sciences. 266:1033–1039. doi: 10.1098/rspb.1999.0740.

Whittington E et al. 2017. Contrasting patterns of evolutionary constraint and novelty revealed by comparative sperm proteomic analysis in Lepidoptera. BMC Genomics. 18:931. doi: 10.1186/s12864-017-4293-2.

Whittington E, Zhao Q, Borziak K, Walters JR, Dorus S. 2015. Characterisation of the Manduca sexta sperm proteome: Genetic novelty underlying sperm composition in Lepidoptera. Insect Biochem Mol Biol. 62:183–193. doi: 10.1016/j.ibmb.2015.02.011.

Wolf KW. 1992. Spindle membranes and microtubules are coordinately reduced in apyrene relative to eupyrene spermatocytes of Inachis io (Lepidoptera, Nympahlidae). Journal of submicroscopic cytology and pathology. 24:381–394.

